# Palm-to-finger cortical functional interactions in primary somatosensory cortex: a 7T fMRI study

**DOI:** 10.1101/2020.09.07.286062

**Authors:** Michel Akselrod, Roberto Martuzzi, Wietske van der Zwaag, Olaf Blanke, Andrea Serino

## Abstract

Many studies focused on the cortical representations of fingers, while the palm is relatively neglected despite its importance for hand function. Here, we investigated palm representation (PR) and its interactions with finger representations (FRs) in primary somatosensory cortex (S1). Few studies in humans suggested that PR is located medially with respect to FRs in S1, yet to date, no study directly quantified the somatotopic organization of PR and the five FRs. Importantly, the relationship between the somatotopic organization and the cortical functional interactions between PR and FRs remains largely unexplored. Using 7T fMRI, we mapped PR and the five FRs at the single subject level. First, we analyzed the cortical distance between PR and FRs to determine their somatotopic organization. Results show that the PR was located medially with respect to D5. Second, we tested whether the observed cortical distances would predict palm-finger functional interactions. Using three complementary measures of functional interactions (co-activations, pattern similarity and resting-state connectivity), we show that palm-finger functional interactions were not determined by their somatotopic organization, that is, there was no gradient moving from D5 to D1, except for resting-state connectivity, which was predicted by the somatotopy. Instead, we show that the representational geometry of palm-finger functional interactions reflected the physical structure of the hand. Collectively, our findings suggest that the spatial proximity between topographically organized neuronal populations do not necessarily predicts their functional interactions, rather the structure of the sensory space (e.g. the hand shape) better predicts the observed functional interactions.

## 1. INTRODUCTION

The sensory cortices of the mammalian brain are topographically organized to form structured maps of the represented sensory features. This organizational principle is preserved across species, sensory modalities and individuals (**Kaas, 1997; Uddin and Fawcett, 1988**) and the role of this topographic organization and its relevance for functional brain interactions are important topics for fundamental and clinical research. Due to metabolic and structural constraints, the spatial proximity between topographically organized neuronal populations directly impacts their functional interactions (i.e. spatially close neurons are more likely to form synapses than distant ones) (**van Ooyen et al., 2014; van Pelt et al., 2013**). On the other hand, the statistics of natural stimulation received during everyday life drives the tuning of functional neuronal interactions through activity-dependent plasticity and can reinforce functional interactions between distant neuronal populations (**Buonomano and Merzenich, 1998**). The somatosensory system is a particularly relevant model to study the relationship between topographic organization in neural maps, functional interactions and use-related function. The somatosensory system must support the functional interactions between elements (i.e. body parts) that can move with respect to each other (e.g. the configuration of the five fingers during object manipulation), can directly interact with each other (i.e. self-touch) and need to fulfill many different sensorimotor functions. While the somatosensory representations of fingers have been researched extensively, the palm has been less studied despite its importance for hand function and despite being anatomically connected with all five fingers. The aim of the present study is to investigate the topographical and functional organization of tactile representations of the palm (palm representation, PR) and its functional interactions with the tactile representations of the five fingers (fingers representations, FRs) in the human primary somatosensory cortex, S1.

S1 is somatotopically organized into a cortical map of the contralateral half of the body (**Kaas et al., 1979, Rasmussen and Penfield, 1947**). FRs in S1 appear in a latero-medial sequence (D1 - D2 - D3 - D4 - D5). This organization is consistent across individuals as shown in recent ultra-high field (7T) fMRI studies (**Besle et al., 2014; Martuzzi et al. 2014; Sanchez-Panchuelo et al. 2010, 2012, 2014; Schweisfurth et al., 2011, 2014, 2018; Schweizer et al., 2008; Stringer et al. 2011, 2014**). While the S1 representations of the base of the fingers (i.e. the distal part of the palm) with respect to FRs have been described in a consistent manner (**Blankenburg et al., 2003; Sanchez-Panchuelo et al. 2012, 2014; Schweisfurth et al., 2011, 2014**), reports concerning the S1 representations of the proximal part of the palm, which is the focus of the present study, are mixed. Several studies suggested that PR is located medially with respect to FRs in S1 (**Blankenburg et al., 2003; Moore et al., 2000; Rasmussen and Penfield, 1947**). However, other studies, specifically aiming at mapping FRs and PR, failed to detect a clear palm representation and could not determine the palm to finger sequence in human S1, likely because they used neuroimaging without sufficient spatial resolution (MEG) or tactile stimulation protocols non-optimized for mapping PR (**Hashimoto et al., 1999; Sanchez-Panchuelo et al., 2012**). Importantly, none of these studies provided direct quantification of the somatotopic organization of the 5 fingers and the palm.

The present study aimed at mapping bilateral PR and FRs in S1 at the level of single subjects and investigate their somatotopic organization and cortical functional interactions. To this aim, we applied a 7T fMRI mapping procedure that was validated in previous studies (**Akselrod et al., 2017; Martuzzi et al., 2014, 2015; Mehring et al., 2019; Serino et al., 2017**). First, we analyzed the cortical distance between PR and the five FRs to determine the somatotopic sequence in S1. This was done in order to validate the proposed serial somatotopic arrangement D1 - D2 - D3 - D4 - D5 - PALM. Second, in order to evaluate the relationship between somatotopic and functional organizations, we tested to which extent the functional interactions between PR and FRs matched their somatotopic organization. We performed the following analyses to obtain complementary measures of functional interactions between PR and FRs: 1) we measured the degree of co-activation between PR and FRs during tactile stimulation of the palm and of the fingers; 2) we compared the multi-voxel patterns of activity in S1 during tactile stimulation of the palm and of the fingers; 3) we quantified the resting-state functional connectivity between PR and FRs.

Based on previous reports in humans (**Blankenburg et al., 2003; Moore et al., 2000; Rasmussen and Penfield, 1947**), we predicted that PR would be located medially with respect to D5 corresponding to a serial somatotopic arrangement. It is possible that the somatotopic arrangement of PR and FRs affects their functional interactions (i.e. stronger functional interactions between closer representations); if this was the case, a serial somatotopic arrangement would predict that functional interactions should be strongest between PALM-D5 representations, with a further decreasing gradient from D4 to D1 and the weakest interaction between PALM-D1 representations. However, considering that the palm is often recruited concomitantly with the five fingers during most in-hand manipulation activities (**Bullock and Dollar, 2011; Pont et al., 2009**) and that functional interactions between hand representations reflect the natural usage of hands (**Ejaz et al., 2015**), we hypothesize that functional interactions between PR and FRs should not match a potential serial somatotopic arrangement (i.e. the palm would not interact preferentially with D5, then D4, D3, …). To provide further insights into the relationship between hand somatotopic organization and hand functional interactions, we compared the representational geometry of the aforementioned measures (i.e. cortical distances, co-activations, multi-voxel activity patterns and functional connectivity) with various competitive models: two models based on the physical structure of the hand (“Body model” and “Perceived body model”) and two purely conceptual models based on different possible configurations of palm and fingers (“Linear model” and “Circular model”).

## 2. METHODS

### 2.1 Subjects

15 healthy subjects (5 females) aged between 18 and 39 years old (mean ± std: 24.3 ± 5.2 years) participated in the study. One participant was excluded due to excessive head motion during MRI acquisition (up to 5mm of movement in the z-direction). Data from another group of 9 healthy subjects (5 females, aged between 26 and 33 years old) recruited in a previous study (**Mehring et al., 2019**) was used to extract average hand dimensions (see section 2.11).

All participants were right-handed as assessed orally using the Edinburgh Handedness Inventory (**Oldfield, 1971**).

All subjects gave written informed consent, all procedures were approved by the Ethics Committee of the Faculty of Biology and Medicine of the University of Lausanne, and the study was conducted in accordance with the Declaration of Helsinki.

### 2.2 Experimental procedure

During fMRI acquisition, subjects received tactile stimulation on six skin regions on both hands (D1 - D2 - D3 - D4 - D5 - PALM). Tactile stimulation consisted of a gentle manual stroking at a rate of approximately 1 Hz performed by an experimenter with his index finger, who received instructions by means of MR compatible earphones. To reduce the variability of the tactile stimulation across participants and to guarantee that a reliable and constant pressure was exerted, the stroking was always performed by the same experimenter, who received extensive training prior to data acquisition. As shown in previous studies, natural touch induces very reliable BOLD signal responses in S1 and is well suited to study body representations in S1 (**Akselrod et al., 2017; Martuzzi et al., 2014, 2015; Serino et al., 2017; van der Zwaag et al., 2015**). The participant’s fingers were repeatedly stroked on the two distal phalanges (thus preventing contamination with palm stimulation), and the palm was stroked in the center on a portion of skin of comparable size. One fMRI run for each hand was acquired in pseudo-randomized order across participants. Within each run, the six regions of the same hand were stroked in a fixed order (D1 - D3 - D5 - D2 - D4 - PALM) and the sequence was repeated 4 times. Stimulation periods of 20 s were interleaved with periods of 10 s of rest (rest periods with no tactile stimulation). In addition to tactile stimulation runs, resting-state data (5min, eyes closed) and anatomical images were acquired for each participant.

### 2.3 Data acquisition

MR images were acquired using a short-bore head-only 7 Tesla scanner (Siemens Medical, Germany) equipped with a 32-channel Tx/Rx RF-coil (Nova Medical, USA) (**Salomon et al., 2014**). Functional images were acquired using a sinusoidal readout EPI sequence (**Speck et al., 2008**) and comprised of 28 axial slices placed approximately orthogonal to the postcentral gyrus (voxel resolution=1.3×1.3×1.3 mm^3^, TR=2s, TE=27ms, flip angle=75°, matrix size=160×160, FOV=210mm, GRAPPA factor=2). The mapping sequence included 361 volumes for each run and the resting-state sequence included 150 volumes. For the resting-state sequence, cardiac and respiratory signals were acquired. Anatomical images were acquired using an MP2RAGE sequence (**Marques et al., 2010**, resolution=1×1×1mm^3^, TE = 2.63ms, TR = 7.2ms, TI1 = 0.9sec, TI2 = 3.2sec, TR_mprage_ = 5sec). To aid coregistration between the functional and the anatomical images, a whole brain EPI volume was also acquired with the same inclination used in the functional runs (81 slices, resolution=1.3×1.3×1.3 mm^3^, TR=5s, TE=27ms, flip angle=75°, matrix size=160×160, FOV=210mm, GRAPPA factor=2).

### 2.4 Data preprocessing

All images were preprocessed using the SPM8 software (Wellcome Department of Cognitive Neurology, London, UK). Preprocessing of fMRI data included slice timing correction, spatial realignment, and minimal smoothing (FWHM=2mm). Freesurfer (http://surfer.nmr.mgh.harvard.edu/, version 6) was used for surface reconstruction (recon-all), for computing cortical distance along the surface (see section 2.6) and for surface rendering of S1 hand maps of a representative subject (Fig.1C). The MRIcron software was used for visualizing results in 3D space for all subjects (McCausland Center for Brain Imaging, University of South Carolina, US, http://www.mccauslandcenter.sc.edu/mricro/mricron). The Connectome Mapper 3 software was used for anatomical parcellation of each subject’s mp2rage data in native space (**Tourbier et al. 2020**).

**Figure 1:**
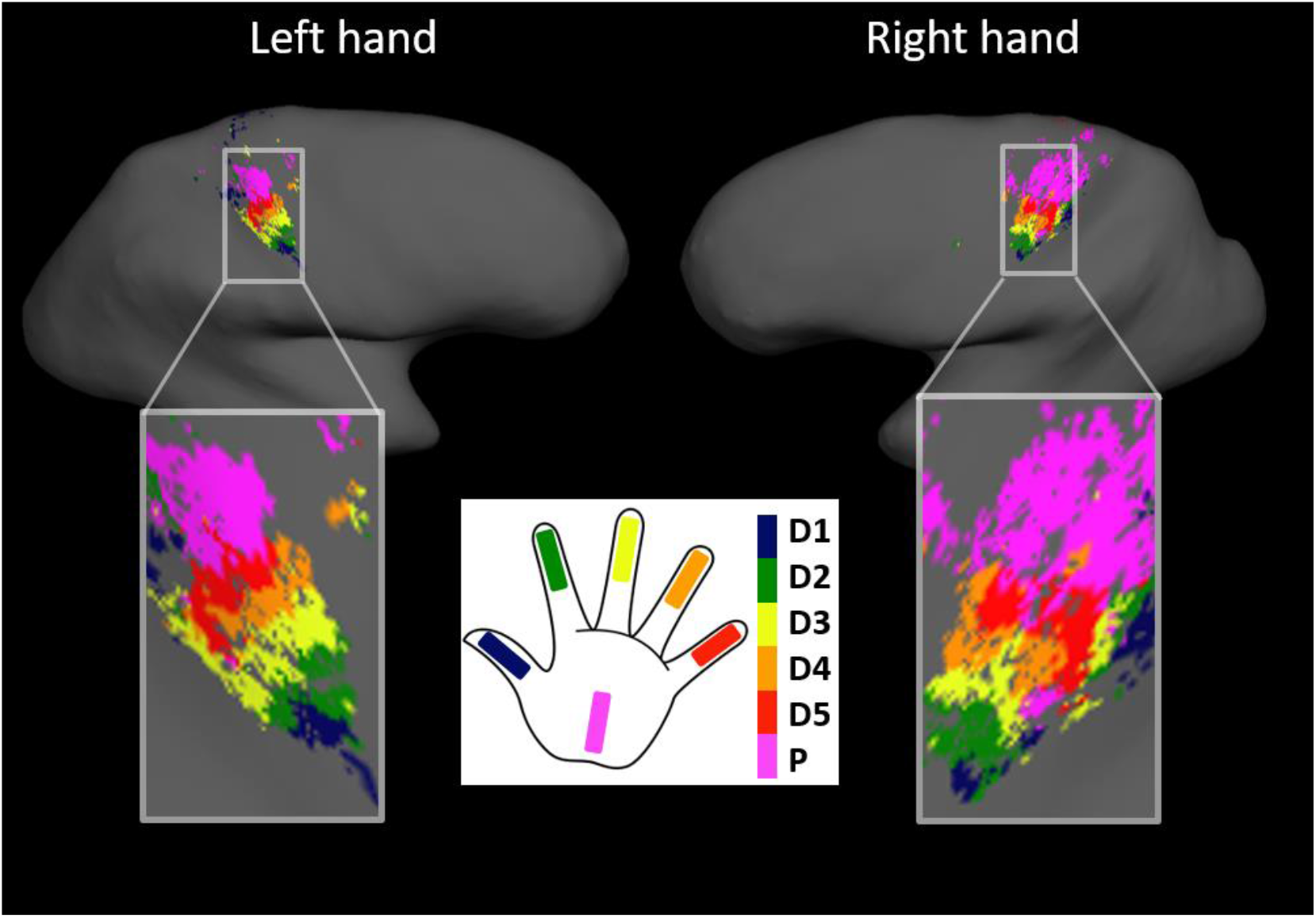
Palm to fingers somatotopy. S1 hand map of a representative subject suggesting the following arrangement in humans: D1 - D2 - D3 - D4 - D5 - PALM.

### 2.5 Definition of somatosensory hand representations

Independently for each subject and each hand, the clusters corresponding to the representations of each stimulated hand region were delimited using an automated approach validated in previous publications (**Akselrod et al., 2017; Martuzzi et al., 2014, 2015; Serino et al., 2017**). A GLM analysis (SPM8) was carried out to estimate the response induced by the stimulation of the different hand regions. The model included 6 regressors (one for each stimulated hand region) convolved with the hemodynamic response and with the corresponding first-order time derivatives, as well as the 6 rigid-body motion parameters as nuisance regressors. For each subject separately, an anatomical parcellation of left and right S1 in native space (i.e. anatomical S1 mask) was computed (Connectome Mapper 3). In addition, for each hand separately, an F-contrast (p,0.0001 uncorrected) across all stimulated hand regions was computed. The active voxels within the F-contrast were used as a functional S1 mask to identify all voxels responding to at least the stimulation of one hand region. Finally, t-contrasts (against rest) were also computed for each stimulated hand region. Then, based on a “winner takes all” approach, each voxel contained within the anatomical and functional S1 masks was labeled as representing the hand region whose stimulation elicited the highest t-score (against rest) for that particular voxel. This approach produces continuous and non-overlapping S1 maps comparable to phase encoding approaches used in mapping studies (**Olman et al. 2010**; **Saadon-Grosman et al. 2015; Sanchez-Panchuelo et al., 2012; Zeharia et al., 2015**).

### 2.6 Analysis of cortical distance

Within each identified hand region representation, the coordinates of the peak activation (maximum t-value) were extracted. These 3D coordinates were transformed into indices of the nearest vertices on the surface space. The surface distances (geodesic) between PR and FRs were calculated for each participant using FreeSurfer (mris_pmake). The statistical analysis described below (section 2.10) was conducted to assess whether the cortical distance was increasing between PR and FRs as expected by a serial somatotopic arrangement: “PALM-D1 > PALM-D2 > PALM-D3 > PALM-D4 > PALM-D5”.

### 2.7 Analysis of co-activations

To investigate the functional interactions between PR and FRs, we computed the co-activations between PR and FRs. To this end, we computed the average BOLD response (beta values) within each FR during the stimulation of the palm (P->FR), as well as the average BOLD response within PR during the stimulation of each finger (F->PR). Co-activations between PR and each FR were defined as the average between P->FR and F->PR. This analysis was conducted in the native space of individual subjects. The statistical analysis described below (section 2.10) was conducted to assess whether co-activations between PR and FRs reflected their somatotopic arrangement.

### 2.8 Analysis of multi-voxel activity patterns

To further investigate the functional interactions between PR and FRs, we compared the multi-voxel activity patterns associated with palm and fingers stimulation (**Kriegeskorte et al, 2008**). Compared to the analysis of co-activations, this measure of functional interactions does not rely on the definition of hand region representations associated with each body part stimulated. Separately for each participant and each hand, we computed a GLM analysis to estimate the beta parameters associated with each period of tactile stimulation (24 tactile stimulation regressors and 6 rigid body motion regressors). Within the active voxels identified to define somatosensory hand representations (see section 2.5), the cross-validated (odd-even split across trials) Mahalanobis distance (**Nili et al. 2014**) between activity patterns associated with palm and fingers stimulation was computed as a measure of pattern dissimilarity. This analysis was conducted in the native space of individual subjects. The statistical analysis described below (section 2.10) was conducted to assess whether the multi-voxel activity patterns associated with palm and fingers stimulation reflected the somatotopic arrangement of PR and FRs.

### 2.9 Analysis of resting-state functional connectivity

Compared to the analyses of co-activations and multi-voxel activity patterns, this measure quantifies functional interactions in the absence of tactile stimulation. Resting-state data were processed using the Conn toolbox (**Withfield-Gabrieli et al., 2012**). At each voxel, the BOLD signal was band-pass filtered (0.008-0.09 Hz). The cardiac and respiratory related components of the BOLD signal were estimated using the RETROICOR algorithm (**Glover et al., 2000**) and regressed out from the data. The average BOLD signal of white matter and cerebrospinal fluid (CSF) and the six estimated motion parameters were also included as nuisance regressors in the model. The bivariate temporal correlations between PR and FRs were calculated from the preprocessed BOLD time-courses of the resting state run. The obtained correlation coefficients were transformed into gaussian values by applying the Fisher transform (**Fisher, 1915**). This analysis was conducted in the native space of individual subjects. The statistical analysis described below (section 2.10) was conducted to assess whether rs-FC between PR and FRs reflected their somatotopic arrangement.

### 2.10 Statistical hypotheses and analyses

First, we used Bayesian statistics to investigate the relationship between PR and FRs across fingers and across body side using the data obtained from the analyses of cortical distance, of co-activations, of multi-voxel activity patterns and of resting-state functional connectivity. Separately for each measure, we computed a two-way Bayesian repeated-measures ANOVA with “FINGER” (5 levels: P-D1, P-D2, P-D3, P-D4 and P-D5) and “SIDE” (2 levels: right hand, RH, and left hand, LH) as within-subject factors (JASP v0.13).

Second, we aimed at validating specific hypotheses regarding the somatotopic and functional organization of hand representations. In particular, we directly tested the hypothesis that PR and FRs are organized in serial arrangement in human S1 (see serial arrangement in Fig.1A, **Moore et al., 2000; Rasmussen and Penfield, 1947**). In addition, we speculated that such organization would not be reflected in the functional interactions between PR and FRs as PR would not interact preferentially with D5, then with D4, then with D3, then with D2 and least with D1 (**Bullock and Dollar, 2011; Pont et al., 2009**). To test these hypotheses, we used the R package *bain* (**Hoijtink et al., 2019**), which allows computing Bayesian statistics based on *informative* hypotheses (i.e. *hypothesis driven* tests). We computed Bayesian one-way repeated-measures ANOVAs separately for each measure (cortical distance, co-activations, multi-voxel activity patterns and resting-state functional connectivity) and each body side (right hand and left hand). We compared three hypotheses: 1) H1, a hypothesis of equivalence between the tested variables with a difference between pairs of variables smaller than a Cohen’s d of 0.2 (P-D1 ≈ P-D2 ≈ P-D3 ≈ P-D4 ≈ P-D5) (**Sawilowsky, 2009**), 2) H_2_, a hypothesis of ordering between the tested variables (P-D1 > P-D2 > P-D3 > P-D4 > P-D5 for cortical distance and pattern dissimilarity or P-D1 < P-D2 < P-D3 < P-D4 < P-D5 for co-activations and functional connectivity), 3) and H_u_ (P-D1, P-D2, P-D3, P-D4, P-D5), the alternative unrestricted hypothesis (i.e. the null hypothesis).

Finally, we computed a Bayesian regression between each measure of functional interactions as observed variable (co-activations, multi-voxel activity patterns and resting-state functional connectivity) and cortical distance as predictor variable (JASP v0.13).

For each Bayesian test, we assumed equal prior probabilities and report the Bayes factors (BF) and posterior probabilities (PP). We considered Bayesian factors >3 as positive evidence, >10 as strong evidence, >30 as very strong evidence (**Kass and Raftery, 1995**).

### 2.11 Dissimilarity analysis

To further investigate the functional organization of hand representations, we conducted dissimilarity analysis (**Akselrod et al., 2017; Kriegeskorte et al, 2008**) and compared the representational geometry associated with the computed measures (cortical distances, co-activations, multi-voxel activity patterns and functional connectivity) with three models of hand representation. Separately for each subject and each hand, the four measures of dissimilarity (cortical distances, co-activations, multi-voxel activity patterns and functional connectivity) were computed between all pairs of fingers in addition to between each finger and the palm to form a 6×6 dissimilarity matrix. The computed dissimilarity matrices were compared with: 1) a model based on the physical shape of the hand, termed “Body” model; 2) a model of somatotopy reflecting the serial arrangements of FRs and PR, termed “Linear” model; 3) a control model, termed “Circular” model.

The “cortical distance” dissimilarity was computed as the surface distance between the coordinates of peak activations associated with the stimulated hand regions similarly to the analysis presented in section 2.6.

The “co-activation” dissimilarity was computed as the co-activations between pairs of S1 hand representations similarly to the analysis presented in section 2.7. The co-activations between pairs of S1 hand representations correspond to a measure of similarity, i.e. pairs of S1 hand representations are considered similar if they are reciprocally co-activated when stimulated separately. The 6×6 similarity matrices of co-activations were transformed into 6×6 dissimilarity matrices by subtracting each co-activation value from the diagonal value of the same row (i.e. *b_i,j_* = *c_i,i_* − *c_i,j_*, where *b* represents the similarity matrix of co-activations, *c* the dissimilarity matrix of co-activations, *i* the row indices and *j* the column indices).

The “multi-voxel activity pattern” dissimilarity was computed as the cross-validated mahalanobis distance between the multi-voxel patterns of S1 activations associated with the stimulated hand regions similarly to the analysis presented in section 2.8.

The “functional connectivity” dissimilarity was computed based on the resting-state functional connectivity between pairs of S1 hand representations similarly to the analysis presented in section 2.9. The resting-state functional connectivity is a measure of similarity, and it was transformed into a measure of dissimilarity by subtracting the bivariate correlation to 1 (i.e. 1-correlation).

The “Body” model was computed using an independent dataset including 9 healthy controls from a previous study (**Mehring et al., 2019**). The data consisted in a localization task, where participants reported the perceived location of different parts of their right hand including the tip and 2^nd^ knuckle of each finger, as well as the center of the palm. The real position of these locations were also recorded. Using these data, we computed average pair-wise distances between the fingers (defined as the average location between the tip and the 2^nd^ knuckle) and the center of the palm. This resulted in a single “Body” model (Fig.6A). The “Linear” model was conceived as a regular decrease in similarity between each adjacent element of the matrix with a step of 1 (arbitrary unit), we note that this model corresponds to a serial model of S1 somatotopy with perfect spacing between representations (Fig.6B). The “Circular model” was conceived as a control model reflecting a plausible geometry of hand representations, but not related to the body or to S1 somatotopy with the palm located in the center and the fingers arranged radially around the palm (Fig.6C).

The matrices corresponding to the four measures of dissimilarity were correlated with the matrices corresponding to the three models separately for each participant (upper part of the matrices were treated as data vectors). For each measure, an upper bound limit of maximum correlation (i.e. noise ceiling) was calculated as the correlation between each subject’s dissimilarity matrix and the group average dissimilarity matrix, averaged across subjects. In order to compare the different models, the resulting correlation values were normalized using the Fischer transformation and statistically analyzed using Bayesian paired t-tests between the model with the highest correlation and the other two models (JASP v0.13). We considered Bayesian factors >3 as positive evidence, >10 as strong evidence, >30 as very strong evidence (**Kass and Raftery, 1995**).

For display purposes, we used classical multidimensional scaling (also known as Principal Coordinate Analysis, **Cooper and Seber, 1985**) to represent the models and the dissimilarity measures on a 2D plot.

### 2.12 Data and code availability statement

The final data presented in the Results section (cortical distances, co-activations, multi-voxel patterns, resting-state functional connectivity and dissimilarity analysis) are openly available on the Zenodo platform. Data analyses were carried out using publicly available resources and/or are fully reproducible from the information provided in the Methods section. Raw data will not be shared to guarantee privacy and confidentiality for the participants.

## 3. RESULTS

Twelve hand regions (6 on each hand) were stimulated during fMRI acquisition to map their cortical representations within S1. This led to a total of 36 mapped representations in S1 per subject (see Methods). Visual inspection of individual maps suggested that the palm (i.e. PRs) was located medially with respect to the D5 FR in all participants. The S1 hand maps of a representative subject are shown in Fig.1 and S1 hand maps for all subjects are shown in supplementary materials (Fig.S1-S2).

### 3.1 Cortical distance

We compared the geodesic distance between PR and each FR using Bayesian statistics (see section 2.10). As shown in Fig.2, the distance between PR and each FR is decreasing when moving from P-D1 to P-D5.

**Figure 2.**
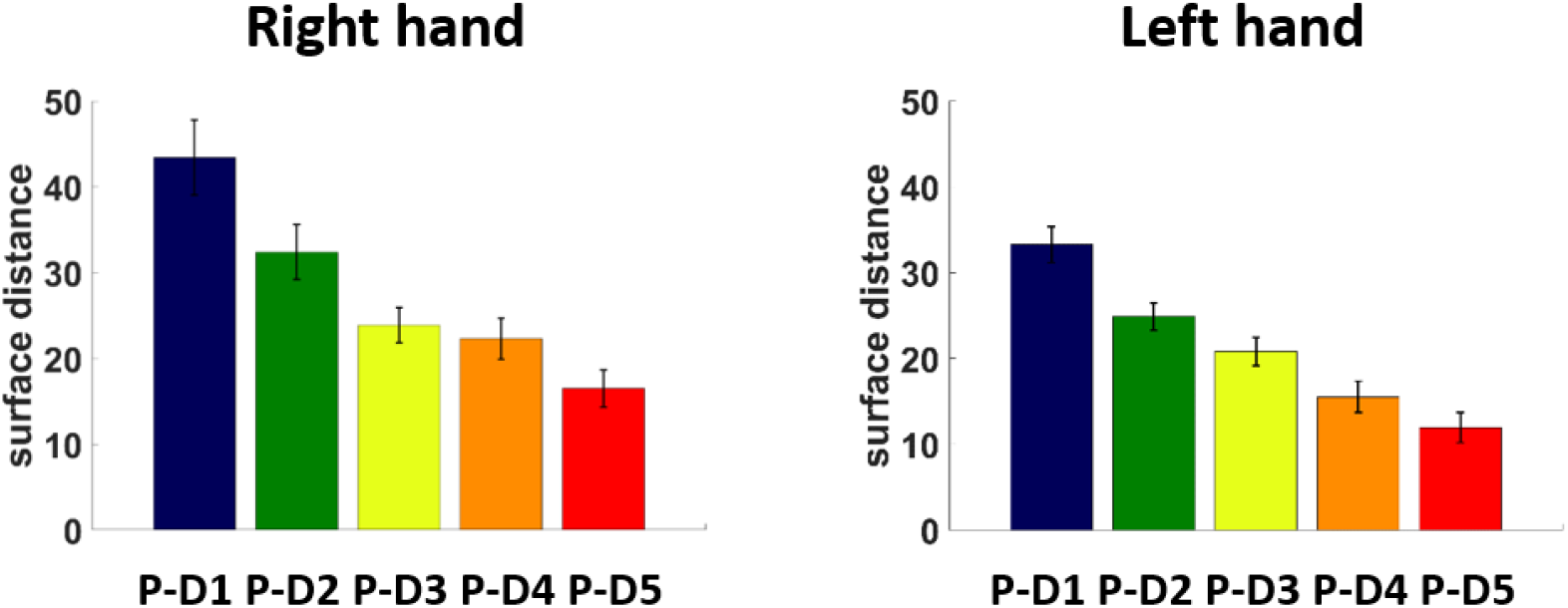
Cortical distance. Bar plots of the cortical distances between PR and each FR in the left hemisphere (right hand representations) and in the right hemisphere (left hand representations). Error bars represent the standard error of the mean.

The two-way ANOVA showed a main effect of finger (BF=2.011e^+18^, PP=0.866), a main effect of body side (BF=3396.92, PP=0.866), but no interaction (BF=0.154, PP=0.134).

The main effect of body side, with very strong evidence (BF>100), was due to reduced distances for left hand representations compared to the right hand.

The *hypothesis driven* ANOVAs strongly supported the ordering hypothesis, *H_2_*, for both hands, suggesting a latero-medial serial arrangement, with the palm located after D5: “D1 - D2 - D3 - D4 - D5 - PALM” (right hand: BF=25.487, PP=0.963; left hand: BF=38.73, PP=0.975, see Tab.1).

**Table 1.**
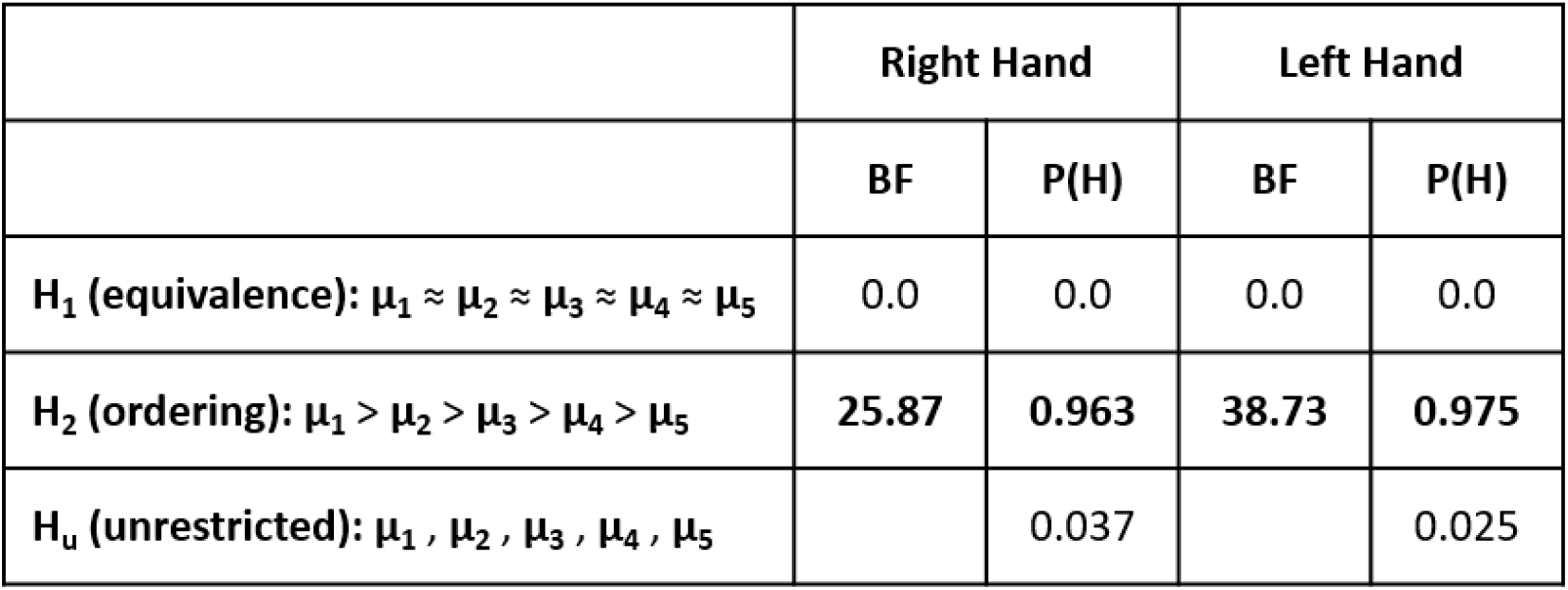
Bayesian statistics on cortical distance. Bayesian ANOVAs were conducted for each hand separately. The Bayes Factor and posterior probability are reported for each tested hypothesis.

|  | Right Hand |  | Left Hand |  |
| --- | --- | --- | --- | --- |
|  | BF | P(H) | BF | P(H) |
| $H_1$ (equivalence): $\mu_1 \approx \mu_2 \approx \mu_3 \approx \mu_4 \approx \mu_5$ | 0.0 | 0.0 | 0.0 | 0.0 |
| $H_2$ (ordering): $\mu_1 > \mu_2 > \mu_3 > \mu_4 > \mu_5$ | 25.87 | 0.963 | 38.73 | 0.975 |
| $H_u$ (unrestricted): $\mu_1, \mu_2, \mu_3, \mu_4, \mu_5$ | | 0.037 | | 0.025 |

To summarize, the analysis of cortical distances comprehensively suggests a serial latero-medial arrangement, with the palm being represented most laterally in human S1. In addition, we found reduced cortical distances between PR and FRs for left hand representations.

### 3.2 Co-activations

We then analyzed the co-activations between PR and FRs, which assess how strongly these representations are reciprocally co-activated when stimulated separately. As shown in Fig.3, there was no consistent evidence of an ordering effect, rather the strongest co-activations are found between P-D1 and P-D5.

**Figure 3.**
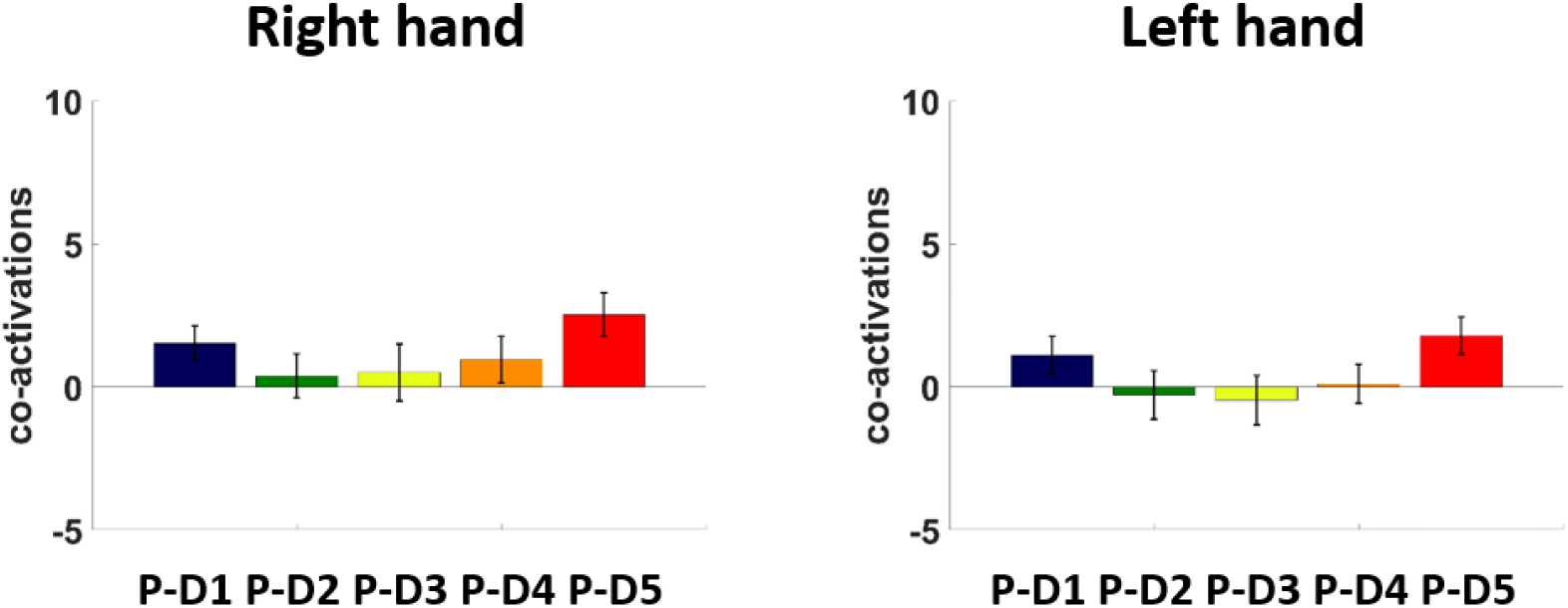
Co-activations. Bar plots of the co-activations between PR and each FR for the right hand (represented in the left hemisphere) and for the eft hand (represented in the right hemisphere). Error bars represent the standard error of the mean.

The two-way ANOVA showed a main effect of finger (BF=12140.48, PP=0.946), a main effect of body side (BF=4.84, PP=0.784), but no interaction (BF=0.069, PP=0.054). The main effect of body side, with positive evidence (BF>3), was due to reduced co-activations for left hand representations compared to the right hand. We note that this is effect is not consistent with the effect of reduced cortical distances for left hand representations, which would rather predict stronger functional interactions with reduced distances.

The *hypothesis driven* ANOVAs supported the unrestricted hypothesis, *H_0_*, for both hands (right hand: PP=0.775; left hand: PP=0.960, see Tab.2). These results show that co-activations between PR and FRs do not follow a pattern predicted by the somatotopic organization and do not show equivalent interactions between PR and the five FRs. Finally, we did not find evidence for a relationship between co-activations and cortical distances (BF=0.484, PP=0.326).

**Table 2.** Bayesian statistics on co-activations. Bayesian ANOVAs were conducted for each hand separately. The Bayes Factor and posterior probability are reported for each tested hypothesis.

|  | Right Hand |  | Left Hand |  |
| --- | --- | --- | --- | --- |
|  | BF | P(H) | BF | P(H) |
| **H<sub>1</sub> (equivalence): $\mu_1 \approx \mu_2 \approx \mu_3 \approx \mu_4 \approx \mu_5$** | 0.0 | 0.0 | 0.0 | 0.0 |
| **H<sub>2</sub> (ordering): $\mu_1 > \mu_2 > \mu_3 > \mu_4 > \mu_5$** | 0.290 | 0.225 | 0.042 | 0.040 |
| **H<sub>u</sub> (unrestricted): $\mu_1, \mu_2, \mu_3, \mu_4, \mu_5$** |  | **0.775** |  | **0.960** |

To summarize, these analyses suggest that functional interactions between PR and FRs, as measured by the degree of mutual co-activations during isolated stimulation, are not equivalent between the PR and each other finger, but rather that that PR might interact preferentially with some FRs, namely D1 and D5. They also do not reflect the somatotopic sequence in S1, suggesting that if a specific pattern of interaction between PR and FRs exist, it does not follow the somatotopic organization. These results might suggest the presence of other patterns of functional interactions that were not formulated in our hypotheses, and therefore, to address this point, we performed dissimilarity analysis that is presented below (3.5).

### 3.3 Multi-voxel activity patterns

We then compared the dissimilarity (i.e. mahalanobis distance) between multi-voxel activity patterns in S1 associated with the tactile stimulation of the palm and of the five fingers. As shown in Fig.5, the highest dissimilarity was observed between P-D3 for both hands, while the lowest dissimilarity was observed between palm-D1 and palm-D5 for both hands. This result is compatible with the co-activation pattern between PR and FRs (i.e. more interaction/similarity between P-D1 and P-D5).

**Figure 4.**
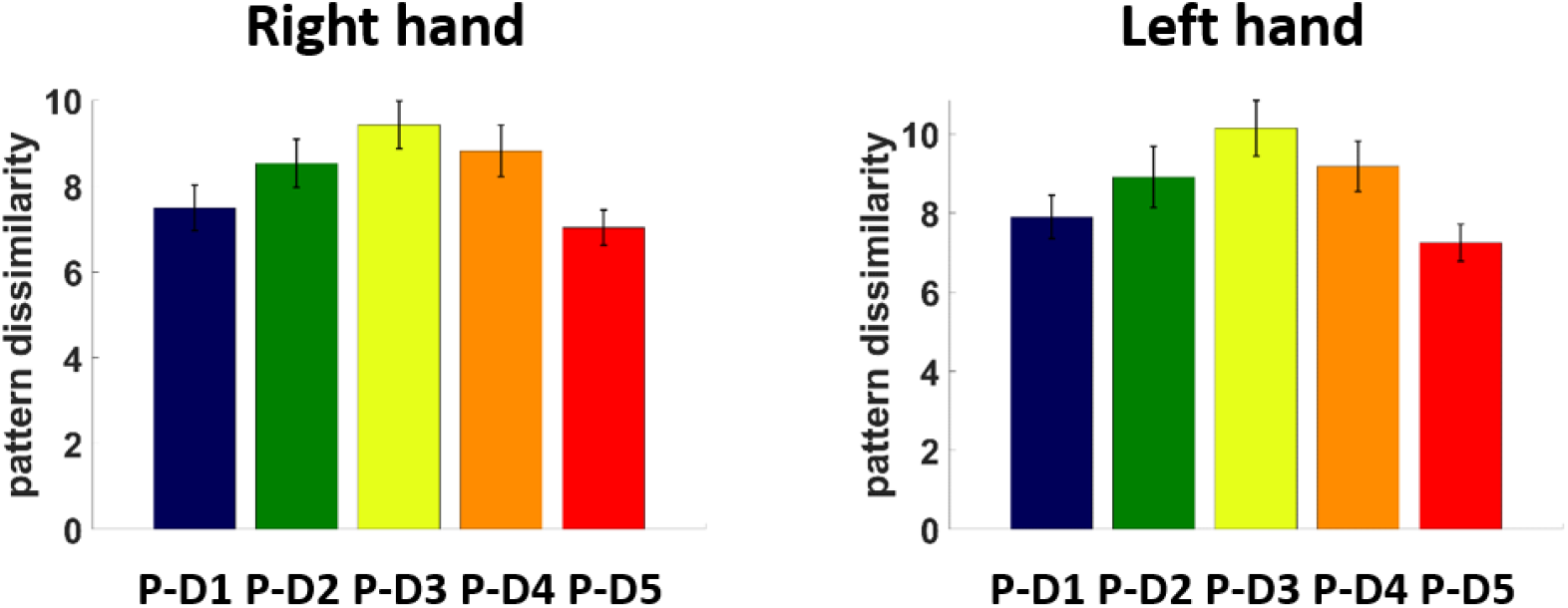
Multi-voxel activity patterns. Bar plots of the dissimilarity between multivoxel activity patterns associated with the stimulation of the palm and of each finger in the left hemisphere (right hand representations) and in the right hemisphere (left hand representations). Error bars represent the standard error of the mean.

**Figure 5.**
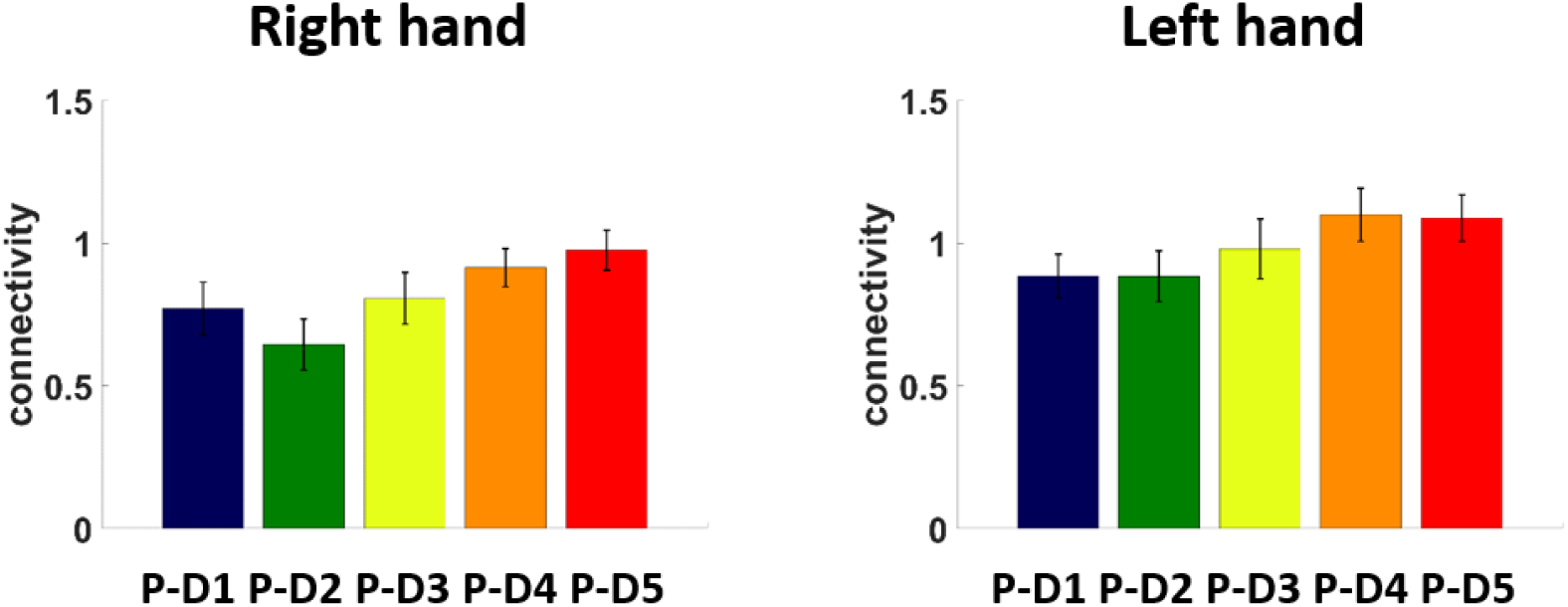
Functional connectivity. Bar plots of the functional connectivity (Z-score) between PR and each of the FR in the left hemisphere (right hand representations) and in the right hemisphere (left hand representations). Error bars represent the standard error of the mean.

The two-way ANOVA showed a main effect of finger (BF=3.796e^+7^, PP=0.973), but no main effect of body side (BF=0.681, PP=0.394) and no interaction (BF=0.069, PP=0.027).

The *hypothesis driven* ANOVAs supported the unrestricted hypothesis, *H_0_*, for both hands (right hand: PP=1.0; left hand: PP=1.0, see Tab.3). These results show that multi-voxel activity patterns do not follow a pattern predicted by the somatotopic organization and do not follow a pattern of equivalence between the palm and the fingers. Finally, we did not find evidence for a relationship between co-activations and cortical distances (BF=0.187, PP=0.158).

**Table 3.**
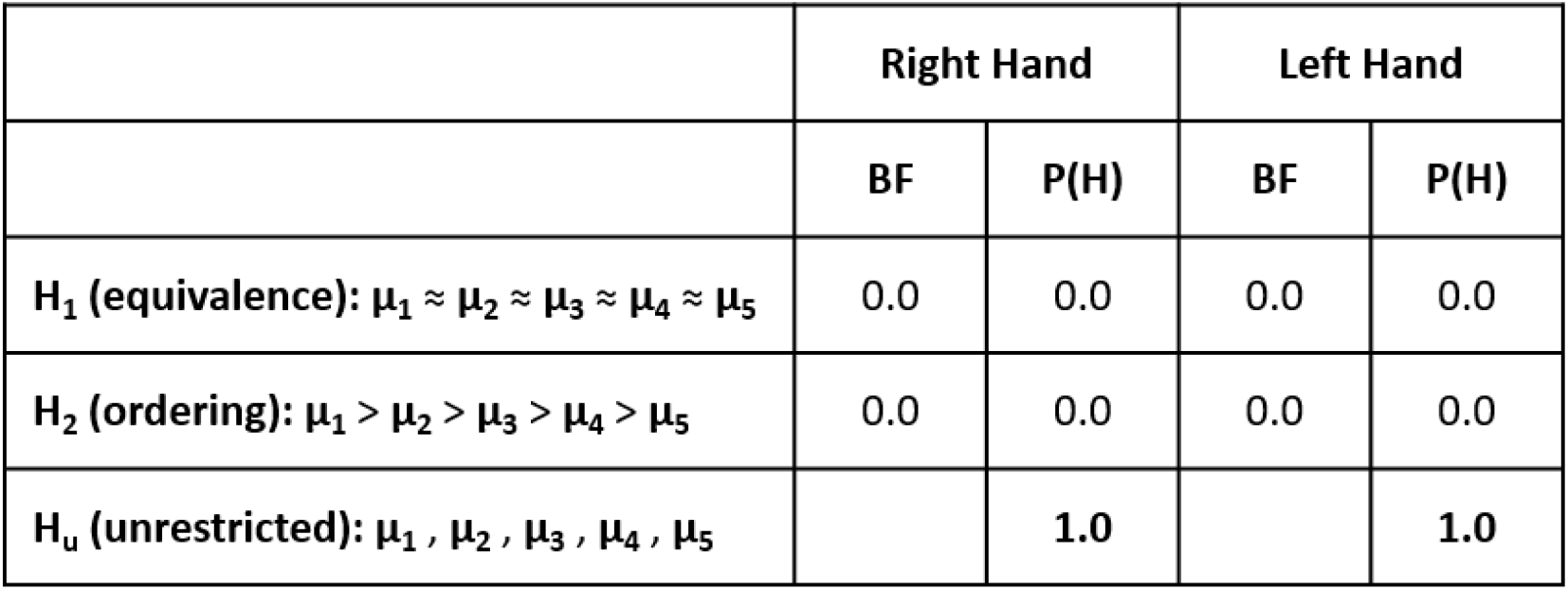
Bayesian statistics on multi-voxel activity patterns. Bayesian ANOVAs were conducted for each hand separately. The Bayes Factor and posterior probability are reported for each tested hypothesis.

To summarize, these analyses suggest that functional interactions between PR and FRs, as measured by multi-voxel activity pattern dissimilarity, do not reflect equivalent interactions between the palm and the five fingers and do not reflect the somatotopic sequence in S1. Consistent with the results of the co-activations analysis (section 3.2), we observed for both hands that minimal pattern dissimilarity was found between P-D1 and P-D5, possibly suggesting the presence of yet another pattern of functional interactions between PR and FRs that was not formulated in our hypotheses (see dissimilarity analysis, 3.5).

### 3.4 Resting-state functional connectivity

Finally, we compared the functional connectivity between PR and FRs. As shown in Fig.6, there is a qualitative trend towards stronger functional connections between PR and FRs which are located closer to PR in S1, although this pattern is not fully consistent (e.g. P-D1 > P-D2 for the right hand, P-D1 ≈ P-D2 and P-D4 ≈ P-D5 for the left hand).

**Figure 6.**
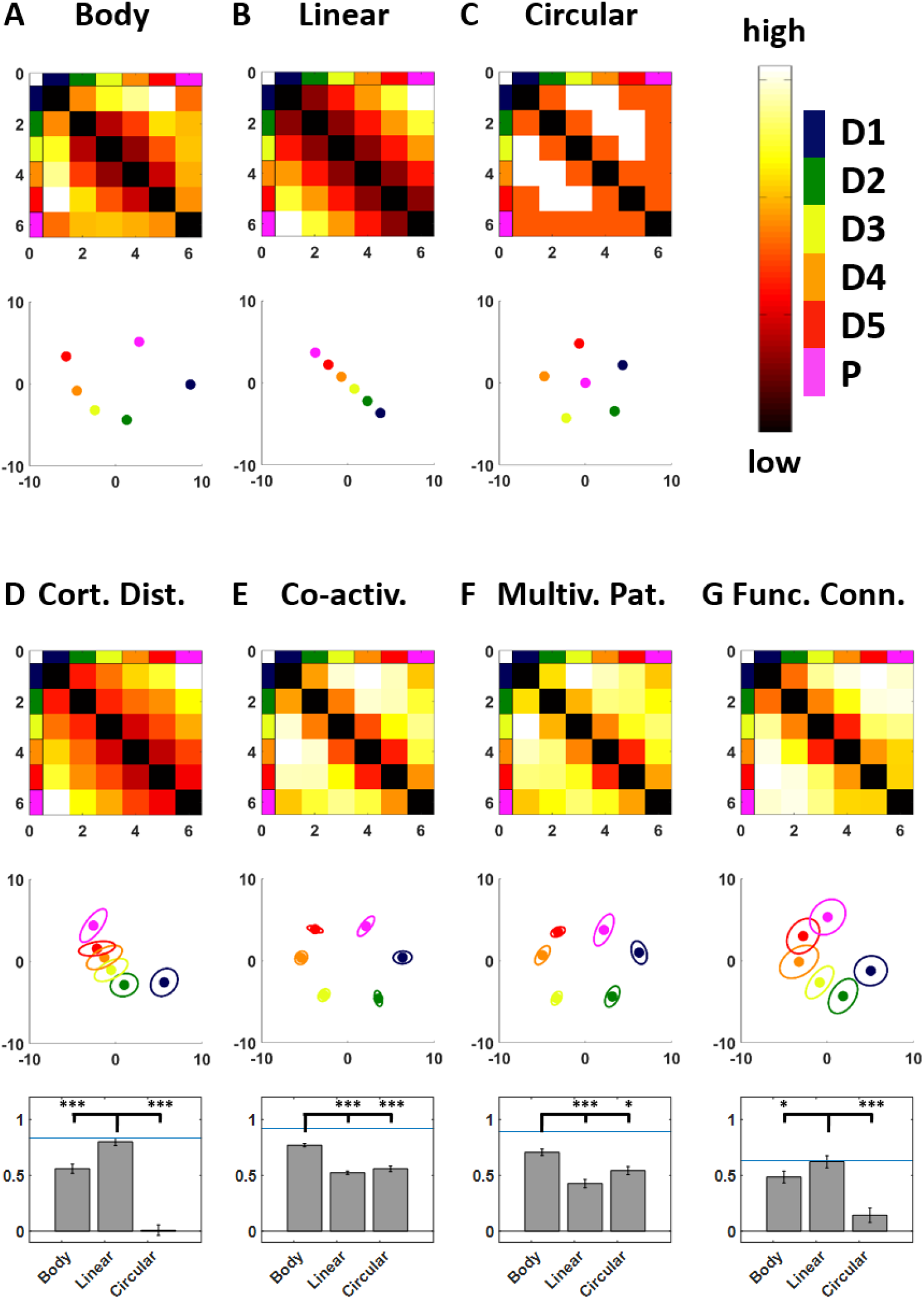
Dissimilarity analysis. A-C. Dissimilarity matrix and 2D configuration for the “Body”, “Linear” and “Circular” models. D-G. Dissimilarity matrix and 2D configuration for the dissimilarity measures based on cortical distances, coactivations, multi-voxel activity patterns and functional connectivity. The correlations between each dissimilarity measure and the three models are shown in the corresponding bar plots. Asterisks indicate the level of evidence found across Bayesian comparisons. The blue line indicates the noise ceiling. Data presented here are averaged across hands.

The two-way ANOVA showed a main effect of finger (BF=113.943, PP=0.914), a main effect of body side (BF=153.667, PP=0.916), but no interaction (BF=0.086, PP=0.078). The main effect of body side, with very strong evidence (BF>100), was due to reduced functional connectivity for right hand representations compared to the left hand. We note that this is effect is consistent with the effect of reduced cortical distances for left hand representations, which would predict stronger functional interactions with reduced distances.

The *hypothesis driven* ANOVAs supported the ordering hypothesis, *H_2_*, for both hands (right hand: BF=1.235, PP=0.553; left hand: BF=44.70, PP=0.978, see Tab.4). We note that, for the right hand, low positive evidence was found for the ordering hypothesis, *H^2^*. These results show that the functional connectivity between PR and FRs follows, at least to some extent, a pattern predicted by the somatotopic organization.

**Table 4.** Bayesian statistics on functional connectivity. Bayesian ANOVAs were conducted for each hand separately. The Bayes Factor and posterior probability are reported for each tested hypothesis.

|  | Right Hand |  | Left Hand |  |
| --- | --- | --- | --- | --- |
|  | BF | P(H) | BF | P(H) |
| $H_1$ (equivalence): $\mu_1 \approx \mu_2 \approx \mu_3 \approx \mu_4 \approx \mu_5$ | 0.0 | 0.0 | 0.0 | 0.0 |
| $H_2$ (ordering): $\mu_1 > \mu_2 > \mu_3 > \mu_4 > \mu_5$ | 1.235 | 0.553 | 44.70 | 0.978 |
| $H_u$ (unrestricted): $\mu_1, \mu_2, \mu_3, \mu_4, \mu_5$ | | 0.447 | | 0.022 |

Finally, we found very strong evidence for a relationship between functional connectivity and cortical distances with stronger functional connectivity being associated with reduced cortical distance (BF=1058.725, PP=0.999).

To summarize, these functional connectivity results suggest that functional interactions between PR and FRs, as measured by resting-state functional connectivity, reflect, at least partially, the somatotopic sequence in S1. This is further supported by the strong evidence for a negative relationship between functional connectivity and cortical distance.

### 3.5 Dissimilarity analysis

Considering that, with the exception of functional connectivity, the measure of functional interactions between PR and FRs were not associated with the somatotopic ordering hypothesis nor with the equivalence hypothesis, we extended previous analyses to investigate the representational geometry of PR and FRs associated with each of the computed measures (cortical distances, co-activations, multi-voxel activity patterns and functional connectivity). We computed dissimilarity matrices based on these four measures and compared them with three models of hand representation, the “Body” model, the “Linear” model and the “Circular” model. Fig.6 shows the four models (A-C) and the three dissimilarity measures (D-G) with their corresponding 2D configuration computed with multidimensional scaling. We note that in this analysis, similar results were obtained for both hands, thus the data were averaged across both hands. Separate data for right and left hands are shown in supplementary materials (Fig.S3-S4).

The three models were designed to capture different aspects of what S1 could represent. The “Body” model formed a 2D configuration compatible with the shape of a hand. The “Linear” model formed a 2D configuration compatible with the somatotopic sequence “D1-D2-D3-D4-D5-PALM”. Finally, the “Circular” model formed a 2D configuration corresponding to a plausible geometry of hand representations, but different from the real shape of a hand and different from S1 hand somatotopic organization.

To assess which model best described the four dissimilarity measures (cortical distances, co-activations, multi-voxel activity patterns and functional connectivity), we computed the correlation between each dissimilarity matrix and the three models and computed Bayesian paired t-tests across these correlations to identify the best models (Tab.S1). For cortical distance, we found that the “Linear” model was the best (r=0.80±0.13) and outperformed the other models with very strong evidence (Linear ≠ Body: BF=173.96, Linear ≠ Circular: BF = 22670.11). For co-activations, we found that the “Body” model was the best (r=0.77±0.06) and outperformed with very strong evidence the “Linear” and “Circular” models (Body ≠ Linear: BF=2.350e^+6^, Body ≠ Circular: BF = 7714.05). For multi-voxel activity patterns, we found that the “Body” model was the best (r=0.71±0.11), outperformed the “Linear” model with very strong evidence (Body ≠ Linear: BF=16619.11) and outperformed with positive evidence the “Circular” model (Body ≠ Circular: BF = 3.77). Finally, for functional connectivity, we found that the “Linear” model was the best (r=0.62±0.20), outperformed the “Body” model with positive evidence (Linear ≠ Body: BF=4.67) and outperformed to “Circular” model with very strong evidence (Linear ≠ Circular: BF = 73.00).

As a control analysis, to confirm that the aforementioned results cannot be explained by the variance associated with the fingers only, we replicated the whole dissimilarity analysis by excluding the palm from the data, leading to 5×5 dissimilarity matrices across the 5 fingers. First, we found that the variance explained by the best models was similar when considering the palm and the five fingers or when considering only the five fingers (Fig.S5). However, when only considering the five fingers the analysis could not disambiguate between the “Body” and “Linear” models. Thus, only when considering the palm and the fingers together, it is possible to highlight a double dissociation between dissimilarity measures best matching the models related to the shape of a hand (co-activations and multi-voxel activity patterns) and dissimilarity measures best matching the model related to somatotopy (cortical distances and functional connectivity).

To summarize, dissimilarity analysis showed that co-activations and multi-voxel activity patterns were related to the shape of a hand, while cortical distances and functional connectivity were rather related to somatotopy. This shows that the representational geometry of hand functional interactions (with the exception of functional connectivity) matched the physical structure of the hand rather than the somatotopic organization of hand representations.

## 4. DISCUSSION

The present study aimed at investigating PR in human S1 and its interactions with the five FRs, by analyzing (1) the cortical distances between somatotopic representations (cortical distance), (2) how PR and FRs co-activate during tactile stimulation (co-activation), (3) the similarity between activity patterns during tactile stimulation (multi-voxel activity pattern), and (4) how PR and FRs are functionally connected to each other (functional connectivity). During the acquisition of fMRI data at ultra-high field (7T), six hand regions (D1 - D2 - D3 - D4 - D5 - PALM) on each side of the body were stimulated using natural touch in a group of healthy subjects. This allowed us to identify the tactile representations of the stimulated hand regions within S1. First, we demonstrated the serial arrangement of the somatotopic sequence: D1 - D2 - D3 - D4 D5 - PALM. Second, we found that this somatotopic sequence is not reflected in the pattern of functional interactions between PR and FRs, with the exception of functional connectivity (see below). Instead, the representational geometry of functional interactions between hand representations better matches the physical shape of the hand rather than the somatotopic organization of its representations.

### 4.1 Mismatch between S1 hand somatotopy and hand structure in humans

The results obtained from the analysis of cortical distances between PR and FRs confirm that the palm representations in human S1 are located medially with respect to the representations of D5, corresponding to a serial somatotopic arrangement in S1 (**Blankenburg et al., 2003; Moore et al., 2000; Rasmussen and Penfield, 1947**). This layout does not correspond to the radial distribution of fingers along the palm on the body, thus creating a discontinuity between S1 hand somatotopy and the physical structure of the hand in humans. Discontinuities between somatotopy and body structure have been well documented in primates (**Felleman et al., 1983; Kaas et al., 1979; Merzenich et al. 1978; Nelson et al. 1980; Rasmussen and Penfield, 1947; Sur et al., 1982**). In particular, the latero-medial arrangement of fingers (D1 - D2 - D3 D4 - D5) in S1, which is found in all primates, forms a hand-arm discontinuity with the latero-medial arrangement of the rest of the arm (distal to proximal).

Interestingly, different somatotopic layouts of the pads (i.e. distal palm and base of the fingers) and fingers have been observed in S1 across primate species (**Felleman et al., 1983; Merzenich et al., 1978; Nelson et al., 1980; Sur et al., 1982**). In Owl and Squirrel Monkeys, the representations of the pads are included in FRs as their most proximal part (**Merzenich et al., 1978; Sur et al., 1982**). This is also the case in humans (**Blankenburg et al., 2003; Sanchez-Panchuelo et al. 2012, 2014; Schweisfurth et al., 2011, 2014)**. Contrastingly, in Cebus and Macaque Monkeys, the representations of the pads lie medially to D5 (**Felleman et al., 1983; Nelson et al., 1980**). Thus, the location of the separation forming the somatotopic hand-arm discontinuity appears to vary across primate species. Furthermore, this somatotopic polymorphism does not correspond to phylogenetic relations between primate species (**Springer et al., 2012**), possibly indicating that a conversion of hand somatotopic layout may have occurred several times during primate evolution. Based on results from this study and previous studies in primates, the proximal part of the palm is represented medially with respect to the 5 fingers in S1 in all studied primate species (present study and **Blankenburg et al., 2003; Felleman et al., 1983; Merzenich et al., 1978; Moore et al., 2000; Nelson et al., 1980; Rasmussen and Penfield, 1947; Sur et al., 1982**). Thus, the proximal part of the palm can be considered a reliable landmark in S1 to study somatotopy within and across primate species.

### 4.2 Mismatch between S1 hand somatotopy and S1 hand functional interactions in humans

Previous studies focusing on fingers reported that S1 functional interactions between FRs, as measured by co-activations, followed a pattern compatible with S1 somatotopy, i.e. the adjacency between FRs in S1 predicts the degree of co-activations (**Besle et al., 2014; Martuzzi et al., 2014**). Similarly, a study focusing on motor representations of fingers showed that multi-voxel activity patterns in S1 associated with finger movements are well described by a somatotopic model of finger adjacency (**Ejaz et. al, 2015**), although these patterns were best described by the natural statistics of hand usage. This suggests that when considering FRs only, a consistency is found between the finger sequence on the hand, the finger somatotopy in S1, and finger functional interactions in S1.

Our analyses of functional interactions between PR and FRs tested whether the S1 palm-to-fingers somatotopy predicts palm-to-fingers functional interactions, as measured by co-activations, multi-voxel activity patterns and resting-state functional connectivity. Importantly, considering the natural usage of the palm in synergy with the fingers for hand function, there is no a priori reason to expect stronger interactions between PR and FRs located closer to PR in S1 (e.g., between palm and D5). Concerning resting-state functional connectivity, dissimilarity analysis showed that functional connectivity matched better with the model reflecting S1 somatotopy. This result can be explained by the well-documented influence of cortical distance on resting-state functional connectivity in both topographically and non-topographically organized brain areas (**Alexander-Bloch et al., 2013; Ercsey-Ravasz et al., 2013; Raemakers et al., 2014; Salvador et al., 2005**). This suggests a possible confound resulting from the use of resting-state functional connectivity to investigate the relationship between functional interactions and cortical topography. More interestingly, our statistical analyses of co-activations and multi-voxel activity patterns revealed that functional interactions between PR and FRs, indeed, do not reflect S1 palm-to-fingers somatotopy. Our analyses of co-activations highlighted that the palm interacts most strongly with D1 and D5. Similarly, multi-voxel activity patterns suggested that palm stimulation induced activity patterns most similar to D1 and D5 stimulation. This pattern of functional interactions is compatible with the physical shape of the hand where, at rest, the tips of D1 and D5 are closer to the center of the palm compared to other fingers. This view was further supported by dissimilarity analysis showing that the representational geometry of hand representations in S1 matched better with the models reflecting the shape of a hand (except for functional connectivity, see below). Another possibility is that the natural statistics of tactile experience during daily life leads to increased likelihood of palm-D1 and palm-D5 co-stimulation (**Ejaz et. al, 2015**). Whether the observed associations (palm-D1 and palm-D5) are better explained by the statistics of use-related tactile stimulation on the hand rather than simply the physical shape of the hand remains to be investigated. Note, however, that natural statistics of tactile stimulation depends on the hand structure, which would make the two hypotheses complementary.

### 4.3 Differences between the right and left hands

Our data also revealed interesting differences between the dominant right hand and the non-dominant left hand in our right-handed participants. We found that the right-hand has overall larger distances between PR and FRs, which might suggest larger cortical territories in S1 for the dominant right hand. However, the inter-digit distances did not differ between right and left hands (Fig.S6), thus suggesting that the aforementioned effect is rather due to PR being located further away from FRs. This is in line with previous fMRI reports showing no difference in size between right and left finger representations (**Boakye et al., 2000; Schweisfurth et al., 2018**). Second, we found overall stronger co-activations between PR and FRs for the right hand compared to the left hand. This effect was not due to simple increased activations, as we found similar strength of activations (within representations) when comparing right and left hand representations (Fig.S7), which corroborates findings from previous fMRI studies showing no difference in strength of activations between right and left hands (**Boakye et al., 2000; Schweisfurth et al., 2018;**but see **Jung et al., 2003, 2008**). Rather, this points towards increased integrative properties between PR and FRs during tactile stimulation. Finally, we also found reduced functional connectivity between PR and FRs for the right hand compared to the left hand, which might be interpreted as increased independence in the absence of stimulation. While this seems counterintuitive with respect to the previous result (increased co-activations during stimulation), this might also suggest a context dependent tuning of integrative properties for the dominant right hand (**Di et al., 2013; Morgan and Price, 2004**).

Results from the present and previous studies (**Boakye et al., 2000; Schweisfurth et al., 2018**) suggest overall no difference between right and left hand representations regarding basic functional properties (extent or strength of activations). This is compatible with behavioural observations showing no difference between right and left hands in tactile spatial acuity (**Sathian and Zangaladze, 1996**). However, our data suggest the presence of differences between the representaiton of the right and the left hand related to integrative properties of somatosensory processing, which might be linked to differences in dexterity associated with hand dominance (**Andersen and Siebner, 2018**).

### 4.4 Plasticity in topographically organized sensory areas

It is believed that topographically organized cortical sensory maps evolved as an optimal solution for energy-efficient spatio-temporal computations (**Kaas, 1997**). It is currently accepted that topographic maps are shaped by a combination of at least two different factors. First, during development, the axonal pathways from the skin to the cortex are established through molecular matching interactions, which is governed by genetics (**Udin and Fawcett, 1988**). The formation of a prototypic topography of sensory maps during development would explain why individuals from a same species share a common architecture. Second, during daily life experience, the spatio-temporal receptive fields of neuronal populations are tuned by sensory stimulation and associated synaptic plasticity (**Buonomano and Merzenich, 1998**).

Our results provide an important account of mismatch between functional interactions and topographical organization. This supports the view that functional cortical interactions, that are consistent with the peripheral structure of the sensory space, can emerge despite the mismatch between topographical organization and the structure of the sensory space. However, our data could not disambiguate between possible contributions of the structure of the sensory space (i.e. the shape of the hand) and of the natural statistics of tactile stimulation received during everyday life (**Ejaz et al., 2015**) in shaping the functional interactions between hand representations. Nevertheless, these two factors are by definition impossible to disentangle in normal conditions, because the physical structure of the body directly impacts the pattern of natural stimulation during everyday life interactions.

Crucially, topographic maps require the continuous competition and interaction between inputs to maintain a normal organization, which is naturally provided during activities of daily living (**Buonomano and Merzenich, 1998**). An extreme example of plasticity in adult primary somatosensory areas is observed following limb amputation (**Flor et al., 1995; Kaas et al., 1983; Makin et al., 2013; Makin and Flor, 2020; Serino et al., 2017**) or to a lesser extent following limb immobilization (**Langer et al., 2012; Liepert et al., 1995; Zannette et al., 1997**). Furthermore, it has been shown that even a brief exposition to repeated sensory stimulation can induce plasticity in primary somatosensory areas (**Godde et al., 1996; Muret et al., 2016; Pleger et al., 2001, 2003**). Similar observations are found for the visual system and the auditory system (**Bilecen et al., 2000; Chino et al., 1992; Kaas et al., 1990; Kaas, 1991; Merabet and Pascual-Lenoe, 2010; Syka et al., 2002**). Considering the results of the present study in light of the capacity of sensory areas to adapt to changes in the structure of the sensory space, a consistency between somatotopic organization and functional interactions could be expected. A possible explanation is that a certain degree of flexibility in the consistency between topographic organization and functional interactions in sensory areas is tolerated. In other words, the metabolic energy cost to tolerate such mismatch is lower than the energy cost required for reorganization. This might suggest that processing mismatched sensory inputs would lead to reorganization only in case of substantial inconsistency.

### 4.4 Study Limitations

We provided tactile stimulation by means of manual stroking delivered by a human experimenter, thus introducing inherent variability in the timing, intensity and extent of stimulation. This choice was motivated by previous work from our group, showing that natural touch is able to induce reliable activations in S1 (**Martuzzi et al., 2014; Akselrod et al., 2017; Serino et al., 2017**), and stronger activations compared to mechanical stimulation (**van der Zwaag et al., 2015**). Although the increased variability associated with natural touch might contribute to the increased signal quality, the lack of controllability might have introduced systematic biases towards a specific body part. Thus, we cannot exclude that at least part of the variance explained by our results might be attributed to the lack of controllability of natural touch.

## 5. CONCLUSIONS

The present study characterizes the properties of PR and its relationship with FRs in human S1. In particular, we investigated the relationship between somatotopic organization and functional interactions of hand representations and reported a mismatch between the two with respect to palm-finger functional interactions. To further study the link between functional properties of tactile hand representations, physical structure of the hand and natural statistics of tactile stimulation, fMRI mapping data (as in the present study) should be combined with behavioral data (e.g. hand tracking during object manipulation). This would allow investigating inter-individual differences in tactile perception and motor skills and would allow studying brain-body plasticity in clinical conditions like amputation or stroke.

## ethics approval statement

Research was conducted under the approval of the Ethics Committee of the University of Lausanne and written informed consent was obtained from all experimental subjects.

## SUPPLEMENTARY FIGURES

**Figure S1.**
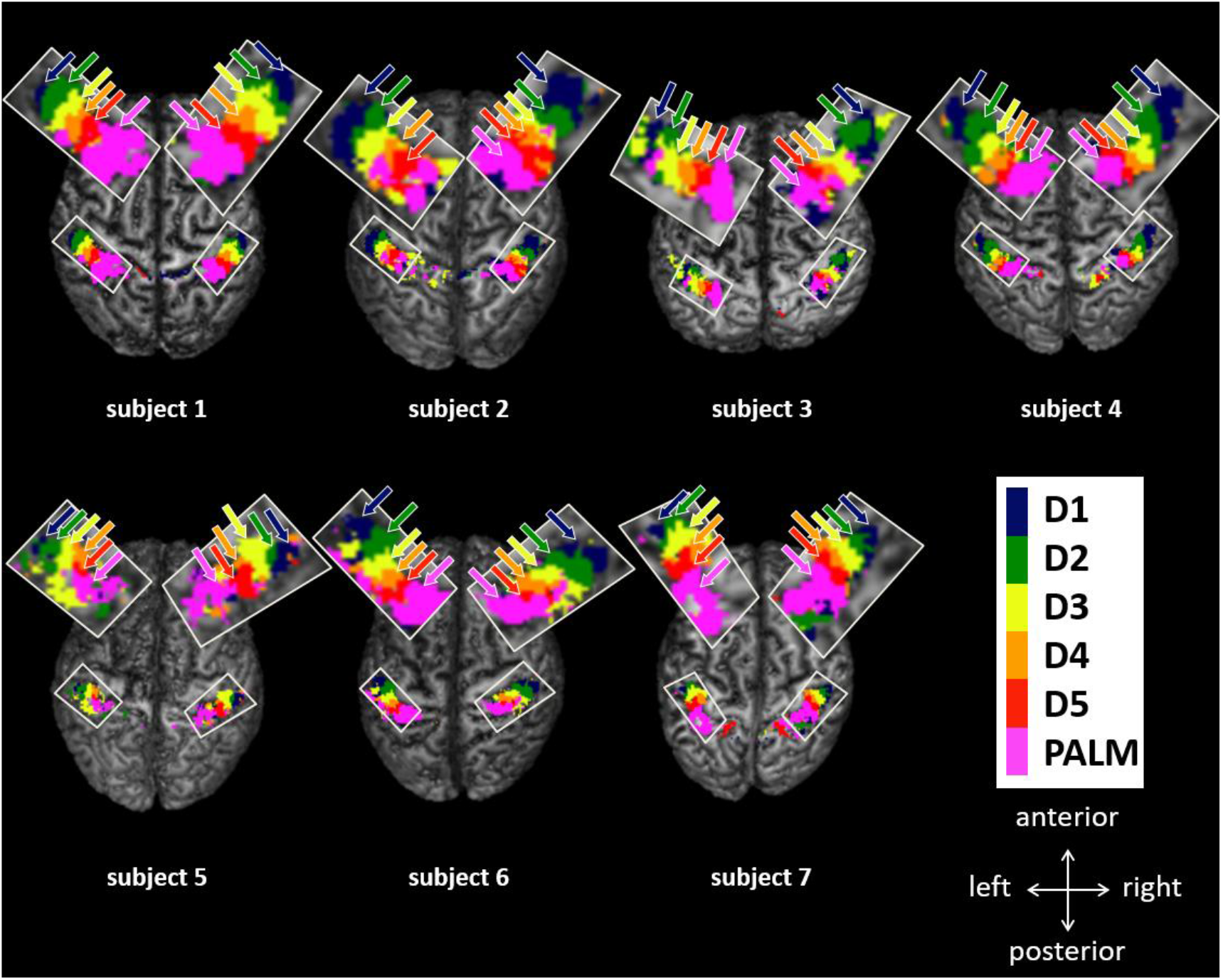
S1 maps from subjects 1-7. S1 hands maps for the right hand in the left hemisphere and the left hand in the right hemisphere are projected and shown from a top view. Note that due to the upward projection used in this visualization, representations located in more superior locations might appear larger as they occlude representations located inferiorly.

**Figure S2.**
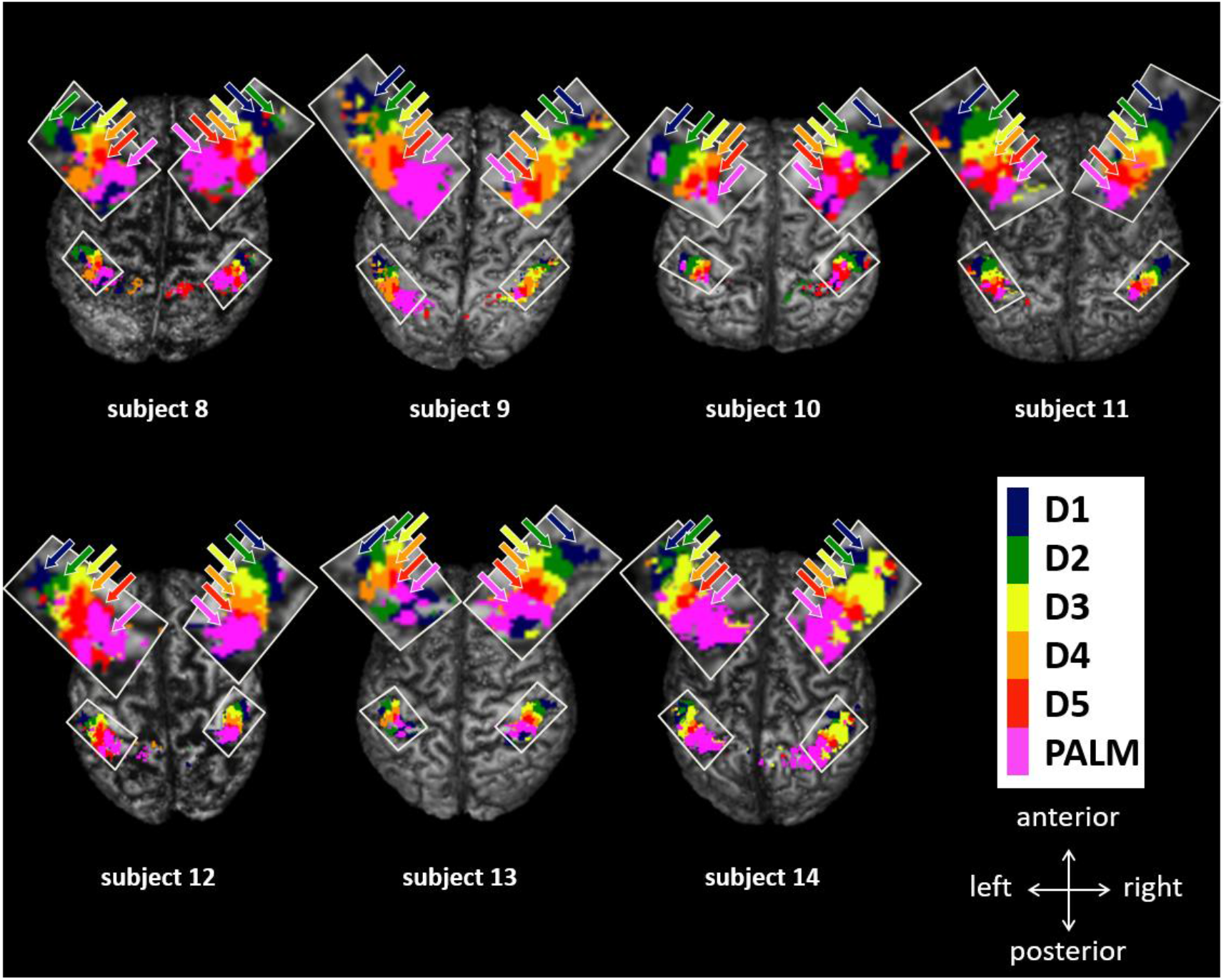
S1 maps from subjects 8-14. S1 hands maps for the right hand in the left hemisphere and the left hand in the right hemisphere are projected and shown from a top view. Note that due to the upward projection used in this visualization, representations located in more superior locations might appear larger as they occlude representations located inferiorly.

**Figure S3.**
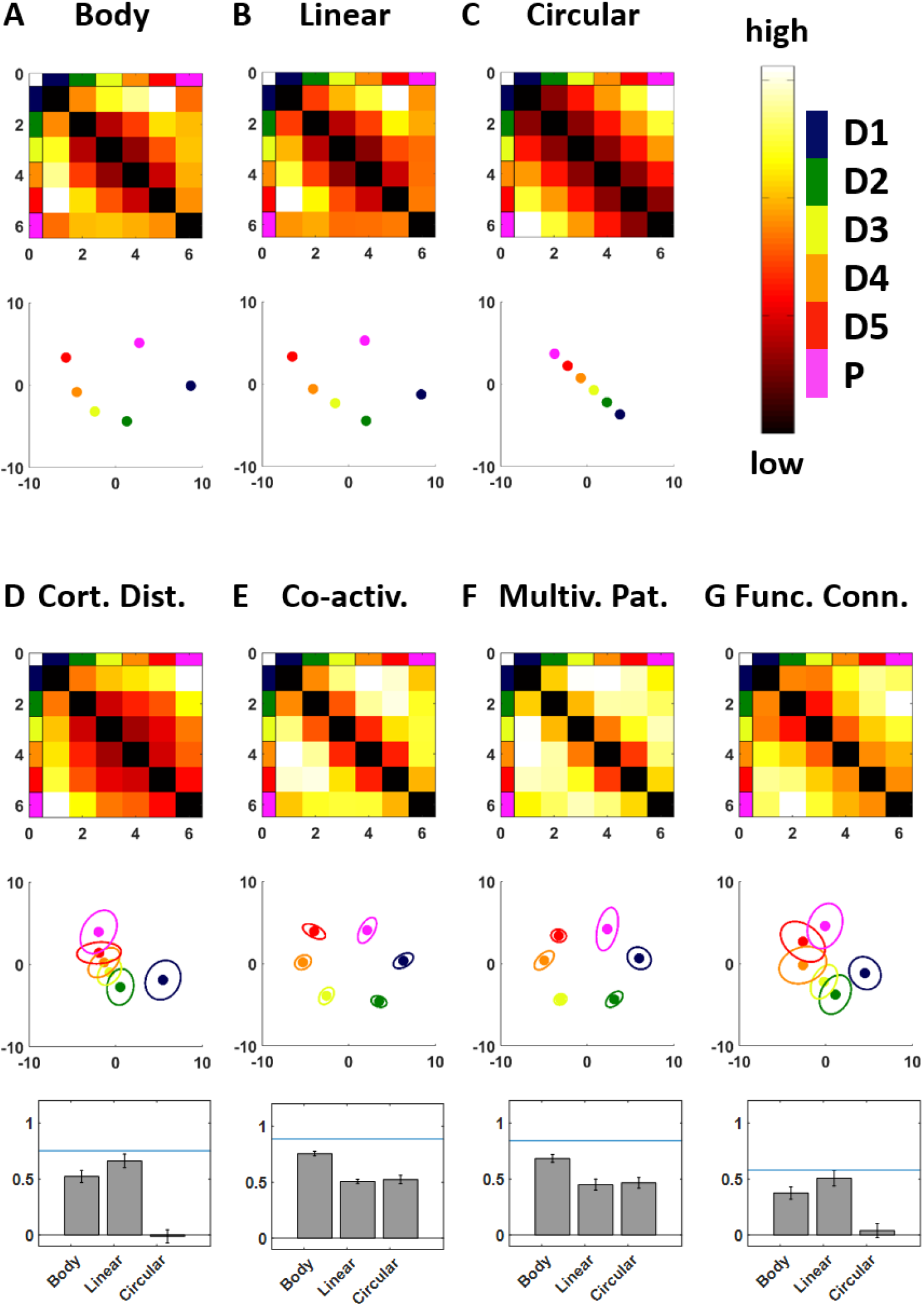
Dissimilarity analysis for the right hand. A-C. Dissimilarity matrix and 2D configuration for the “Body”, “Linear” and “Circular” models. D-G. Dissimilarity matrix and 2D configuration for the dissimilarity measures based on cortical distances, co-activations, multi-voxel activity patterns and functional connectivity. The correlations between each dissimilarity measure and the three models are shown in the corresponding bar plots. Asterisks indicate the level of evidence found across Bayesian comparisons. The blue line indicates the noise ceiling. Data presented here are averaged across hands.

**Figure S4.**
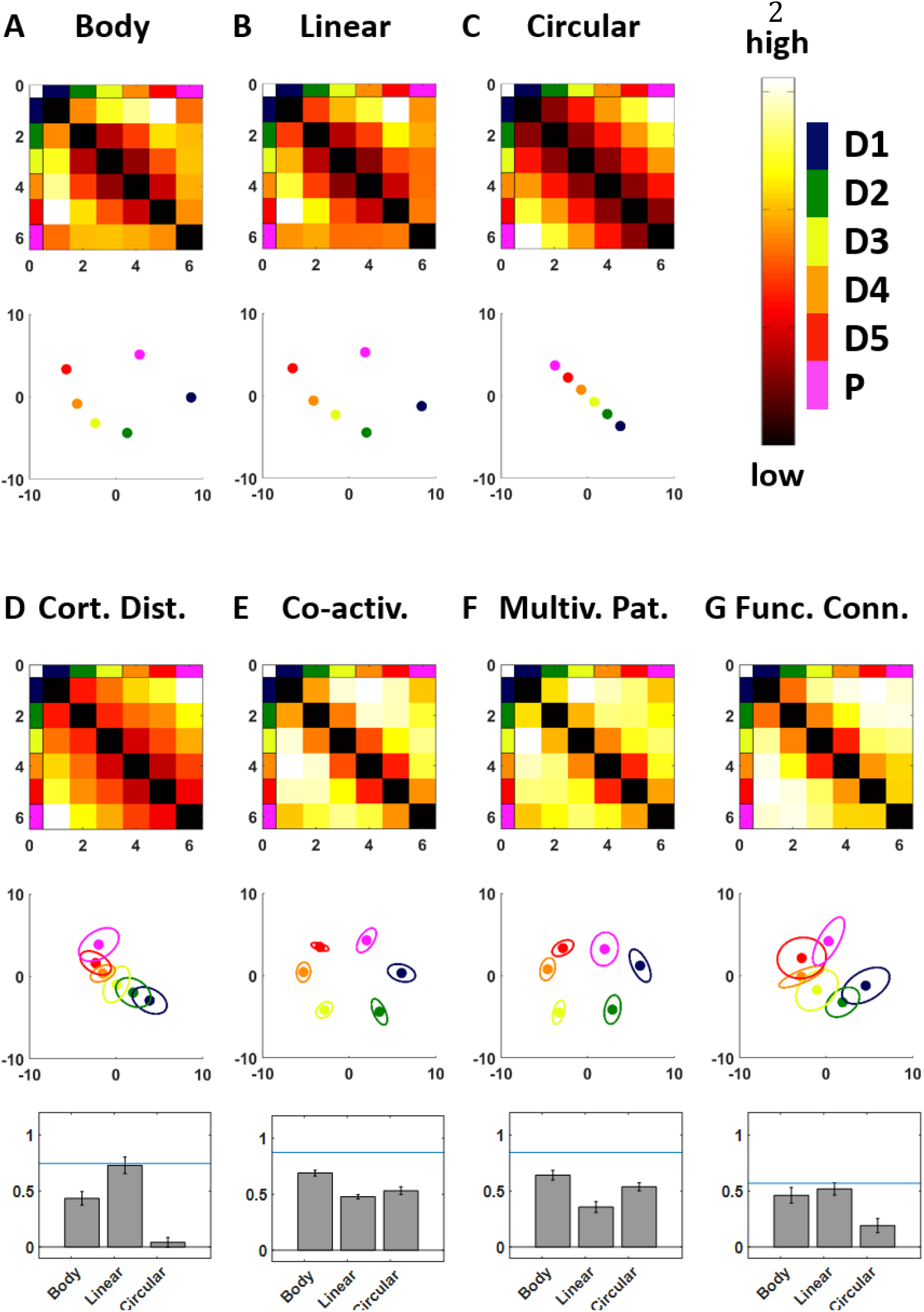
Dissimilarity analysis for the left hand. A-C. Dissimilarity matrix and 2D configuration for the “Body”, “Linear” and “Circular” models. D-G. Dissimilarity matrix and 2D configuration for the dissimilarity measures based on cortical distances, co-activations, multi-voxel activity patterns and functional connectivity. The correlations between each dissimilarity measure and the three models are shown in the corresponding bar plots. Asterisks indicate the level of evidence found across Bayesian comparisons. The blue line indicates the noise ceiling.

**Figure S5.**
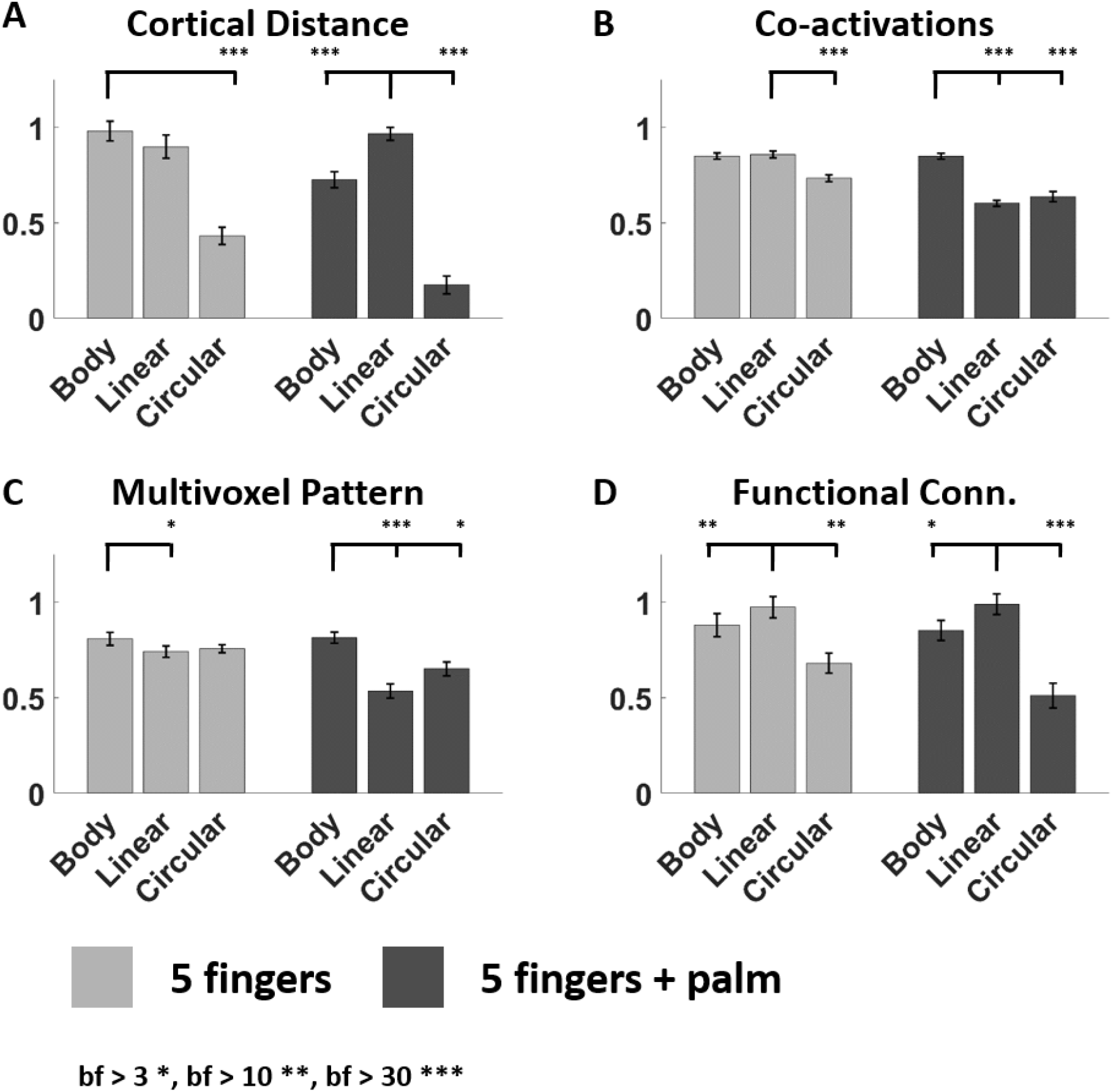
Comparison between dissimilarity analyses. Dissimilarity analysis was replicated to consider the 5 fingers only by excluding data from the palm. Note that in both cases the variance explained is very high for the best models. However only data including the palm is able to highlight the double dissociation showing that the measures of cortical distance and functional connectivity are better described by the linear model, and that the measures of co-activations and multivoxel activity patterns are better described by the body model. Data were normalized with respect to noise ceiling.

**Table S1.** Bayesian comparisons for dissimilarity analysis with the palm and the five fingers. Separately for each measure of dissimilarity (cortical distances, coactivations, multi-voxel activity patterns and functional connectivity), paired comparisons were computed between each pair of models. Comparisons highlighted in bold are reported in Fig.6 and Fig.S5.

| Measure | Comparison | BF |
| --- | --- | --- |
| **Cortical distance** | **Body ≠ Linear** | **173.96** |
| Cortical distance | Body ≠ Circular | 55616.24 |
| **Cortical distance** | **Linear ≠ Circular** | **22670.11** |
| **Co-activation** | **Body ≠ Linear** | **2.350e<sup>+6</sup>** |
| **Co-activation** | **Body ≠ Circular** | **7714.05** |
| Co-activation | Linear ≠ Circular | 0.481 |
| **Multi-voxel patt.** | **Body ≠ Linear** | **16619.11** |
| **Multi-voxel patt.** | **Body ≠ Circular** | **3.77** |
| Multi-voxel patt. | Linear ≠ Circular | 0.90 |
| **Functional Conn** | **Body ≠ Linear** | **4.67** |
| Functional Conn | Body ≠ Circular | 50.23 |
| **Functional Conn** | **Linear ≠ Circular** | **73.00** |

**Table S2.** Bayesian comparisons for dissimilarity analysis with only the five fingers. Separately for each measure of dissimilarity (cortical distances, co-activations, multi-voxel activity patterns and functional connectivity), paired comparisons were computed between each pair of models. Comparisons highlighted in bold are reported in Fig.S5.

| Measure | Comparison | BF |
| --- | --- | --- |
| Cortical distance | Body ≠ Linear | 1.06 |
| **Cortical distance** | **Body ≠ Circular** | **4357.32** |
| Cortical distance | Linear ≠ Circular | 247.32 |
| Co-activation | Body ≠ Linear | 0.34 |
| Co-activation | Body ≠ Circular | 14.86 |
| **Co-activation** | **Linear ≠ Circular** | **18.22** |
| **Multi-voxel patt.** | **Body ≠ Linear** | **4.39** |
| Multi-voxel patt. | Body ≠ Circular | 0.57 |
| Multi-voxel patt. | Linear ≠ Circular | 0.27 |
| **Functional Conn** | **Body ≠ Linear** | **14.67** |
| Functional Conn | Body ≠ Circular | 2.68 |
| **Functional Conn** | **Linear ≠ Circular** | **17.58** |

**Figure S6.**
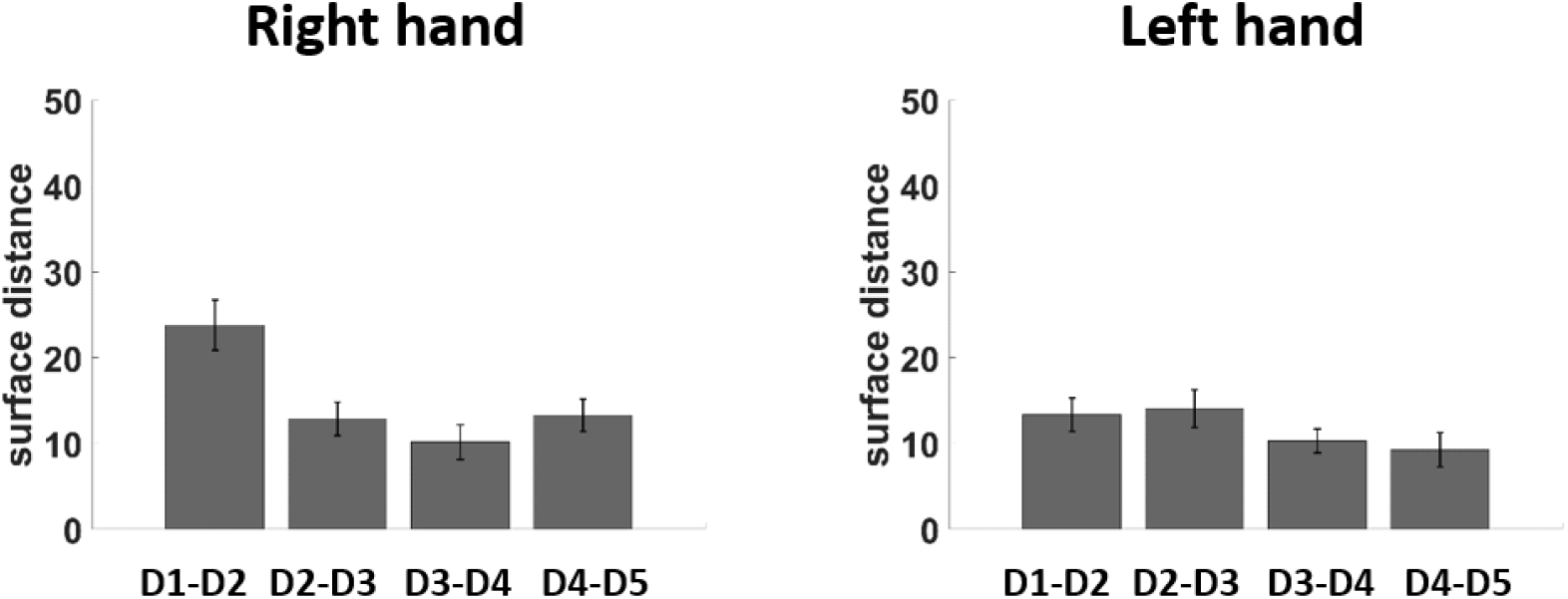
Inter-digit cortical distance. Bar plots of the cortical distances between each pair of FR in the left hemisphere (right hand representations) and in the right hemisphere (left hand representations). Error bars represent the standard error of the mean. A two-way Bayesian ANOVA showed very strong evidence for a main effect of “Finger pair” (BF=108.70, PP=0.246), no evidence for a main effect of side (BF=2.32, PP=0.174) and positive evidence for an interaction (BF=4.36, PP=0.75).

**Figure S7.**
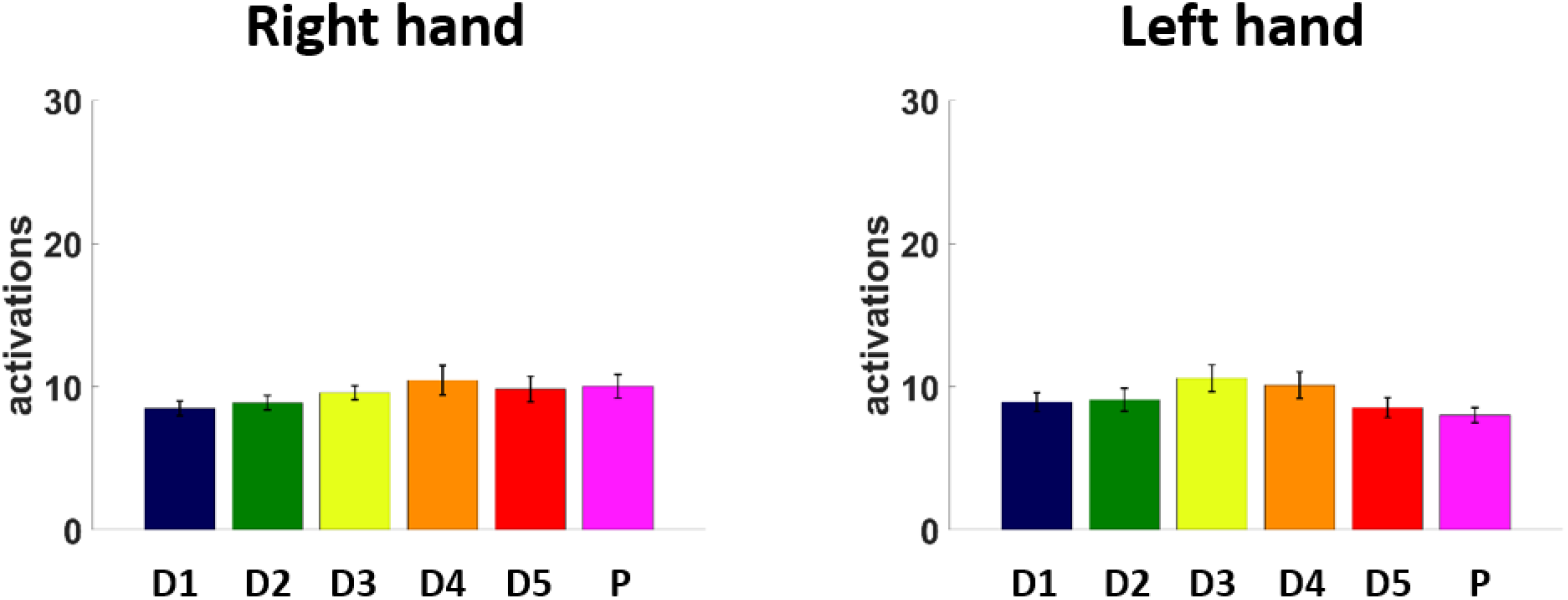
Activations within PR and FRs. Bar plots of beta activations within PR and FRs in the left hemisphere (right hand representations) and in the right hemisphere (left hand representations). Error bars represent the standard error of the mean. A two-way Bayesian ANOVA showed positive evidence for a main effect of “Representation” (BF=3.48, PP=0.566), no evidence for main effect of side (BF=0.296, PP=0.166) and no evidence for an interaction (BF=2.08, PP=0.27).

